# Anacardic acid protects against phenylephrine-induced mouse cardiac hypertrophy through JNK signaling-dependent regulation of histone acetylation

**DOI:** 10.1101/2020.02.06.937672

**Authors:** Bohui Peng, Chang Peng, Xiaomei Luo, Lixin Huang, Qian Mao, Huanting Zhang, Xiao Han

## Abstract

Cardiac hypertrophy is a complex process induced by the activation of multiple signaling pathways. We previously reported that anacardic acid (AA), a histone acetylase (HAT) inhibitor, attenuates phenylephrine (PE)-induced cardiac hypertrophy by downregulating histone H3 acetylation at lysine 9 (H3K9ac). Unfortunately, the upstream signaling events remained unknown. The mitogen-activated protein kinase (MAPK) pathway is an important regulator of cardiac hypertrophy. In this study, we explored the role of JNK/MAPK signaling in cardiac hypertrophy. A mouse model of cardiomyocyte hypertrophy was successfully established *in vitro* using PE. This study showed that p-JNK directly interacts with HATs (P300 and P300/CBP-associated factor, PCAF) and alters H3K9ac. In addition, both the JNK inhibitor SP600125 and the HAT inhibitor AA attenuated p-JNK overexpression and H3K9 hyperacetylation by inhibiting P300 and PCAF during PE-induced cardiomyocyte hypertrophy. Moreover, we demonstrated that both SP600125 and AA attenuate the overexpression of cardiac hypertrophy-related genes (*MEF2A, ANP, BNP*, and *β-MHC*), preventing cardiomyocyte hypertrophy and dysfunction. These results revealed a novel mechanism through which AA might protect mice from PE-induced cardiac hypertrophy. In particular, AA inhibits the effects of JNK signaling on HAT-mediated histone acetylation, and could therefore be used to prevent and treat hypertrophic cardiomyopathy.

## 1. Introduction

Cardiac hypertrophy is an independent predictive factor of cardiovascular diseases and ultimately leads to heart failure and sudden cardiac death [1]. In recent years, cardiac hypertrophy has become one of the main causes of morbidity and mortality in modern society [2]. Unfortunately, currently available treatments can only temporarily stop disease progression. Therefore, increasing efforts are being made to develop more effective preventive and therapeutic measures to cure cardiac hypertrophy. Several studies have shown that cardiac hypertrophy is induced by the activation of multiple signaling pathways, and the mitogen-activated protein kinase (MAPK) signaling pathway is believed to play an essential pathogenetic role [3,4]. It is well known that phenylephrine (PE) is one of the important factors regulating cardiac hypertrophy and its such function is realized through complex intracellular signal transduction pathways as follows: MAPK signaling pathway [5]. JNK is a member of the MAPK protein family and has been previously related to cardiac hypertrophy [6]. It is found that JNK signaling pathway can regulate various physiological processes including cardiomyocyte hypertrophy, apoptosis and inflammation [4]. We previously study demonstrated that a Chinese herbal extract containing anacardic acid (AA) attenuates cardiac hypertrophy by repressing histone acetyltransferase (HAT) activity, thus downregulating histone hyperacetylation [7]. However, the signaling events upstream of this pathway remained unclear. Even more, whether the JNK signaling pathway is directly involved anacardic acid attenuates cardiac hypertrophy have not been elucidated. Herein, The purpose of this study was to determine whether JNK signaling is involved in the AA-mediated attenuation of cardiac hypertrophy.

## 2. Materials and methods

### 2.1. Experimental mice

Sterile or pathogen-free male and female 1-3-day-old Kunming mice with a body mass of 2.3-2.7 g were obtained from the Experimental Animal Center at Chongqing Medical University (Chongqing, China). All experimental procedures were approved by the Animal Care and Use Committee of Zunyi Medical University.

### 2.2. Cell culture

Neonatal 1-3-day-old Kunming mice were sacrificed by decapitation under aseptic conditions, and the hearts were excised immediately. The cardiac ventricles were cut into pieces of approximately 1–2 mm^3^ and dissociated by trituration in 0.05% collagenase type II (Worthington, Lakewood, NJ, USA) 8-10 times for 5 min each. Cells were finally obtained by centrifugation for 10 min at 1500 × *g* and after discarding the supernatant, they were resuspended in DMEM/F12 (1:1) containing 20% fetal bovine serum (Invitrogen, Carlsbad, CA, USA). Subsequently, the cells were incubated for 1.5 h at 37 °C in a humidified atmosphere with 5% CO_2_ to separate out the fibroblasts. The cardiomyocytes were randomly divided into six groups: Control group; Vehicle group, 100 μmol/L PE (Sigma, St. Louis, MO, USA) and isopyknic DMSO (Solarbio, Beijing, China) treatment; PE group, 100 μmol/L PE treatment; AA group, 50 μmol/L AA (Sigma, St. Louis, MO, USA) and 100 μmol/L PE treatment; AA+SP group, 50 μmol/L AA, 20 μmol/L JNK inhibitor SP600125 (Calbiochem, Darmstadt, Germany) and 100 μmol/L PE treatment; SP group, 20 μmol/L JNK inhibitor SP600125 and 100 μmol/L PE treatment.

### 2.3. Cell viability assay

The cardiomyocytes were seeded in a 96-well plate at a density of 2 × 10^4^ cells/well. Next, the cells were added to 10 µl Cell Counting Kit-8 (CCK-8) solution (Solarbio, Beijing, China) and incubated for 4 h at 37 °C in the dark. Finally, the absorbance was measured at 450 nm using a Universal Microplate Spectrophotometer (Bio-Rad, CA, USA).

### 2.4. Western blotting

The cardiomyocytes were collected, and nucleoproteins were extracted using a nuclear protein extraction kit (Merck Millipore, Darmstadt, Germany). Nucleoproteins were separated by electrophoresis on 6/12% sodium dodecyl sulfate (SDS) polyacrylamide gels and blotted onto polyvinylidene difluoride (PVDF) membranes (Merck Millipore, Darmstadt, Germany). After blocking for 1 h with 5% bovine serum albumin, these PVDF blots were probed with rabbit polyclonal antibodies against brain natriuretic peptide (BNP) (Abcam, 1:1000 dilution), atrial natriuretic peptide (ANP) (Abcam,1:5000 dilution), lysine 9-acetylated histone H3 (H3K9ac) (Abcam, 1:5000 dilution), beta-myosin heavy chain (β-MHC) (Abcam, 1:5000 dilution), PCAF (Abcam, 1:500 dilution), and P300 (Abcam, 1:2000 dilution), rabbit polyclonal antibodies against JNK and phospho-JNK (Cell Signaling, 1:1000 dilution), or rabbit polyclonal antibodies against β-actin and histone H3 (Beyotime, 1:5000 dilution). All antibodies were diluted in tris-buffered saline with tween 20 (TBST) containing 5% non-fat milk, and incubations were performed at 4 °C overnight. HR-conjugated goat anti-rabbit antibody (Santa Cruz Biotechnology, Texas, USA, 1:2000 dilution) was used as the secondary antibody. After scanning, the bands were subjected to analysis using the Quantity One (Version 4.4) software package (Bio-Rad, CA, USA).

### 2.5. Total RNA extraction and real-time quantitative polymerase chain reaction (RT-PCR)

Total RNA from cardiomyocytes was extracted with an RNA extraction kit (BioTeke, Beijing, China) following the manufacturer’s protocol. The RNA was reverse-transcribed to single-stranded cDNA using the AMV Reverse Transcription System (Takara, Dalian, Liaoning, China). Then, cDNA was amplified with a SYBR Green dye kit and gene-specific primers (Takara, Shiga, Japan). Data were normalized to β-actin mRNA levels. Relative gene expression was determined by the 2^-ΔΔ Ct^ method.

### 2.6. Immunofluorescence

The cardiomyocytes were seeded in 6-well plates (1 × 10^5^ cells/well). After 24 h of culture, cells were incubated for 1 h with 50 μmol/L AA, followed by the addition of 20 μmol/L SP600125, and for a further 48 h with 100 μmol/L phenylephrine. Then, the cells were fixed with 4% paraformaldehyde at room temperature (RT) for 15 min. Next, the cells were treated with 0.3% triton X-100 in phosphate buffer saline at RT for 20 min, incubated with 10% goat serum at 37 °C for 30 min, and then incubated with primary antibodies against α-actinin (1:100, Proteintech), H3K9ac (1:1000, Abcam), P300 (1:200, Abcam), and PCAF (1:250, Abcam) at 4 °C overnight. Then, the cells were incubated with Alexa Fluor 594 goat anti-mouse IgG secondary antibody (1:1000, Thermo Fisher Scientific) and Alexa Fluor 488 goat anti-rabbit IgG (1:200, Invitrogen) secondary fluorescence antibodies for 1 h at 37 °C in the dark. Finally, the cells were counterstained with DAPI at RT for 5 min. All images were taken under a fluorescence microscopy using the same imaging parameters, and fluorescence quantification was performed using ImageJ software.

### 2.7. Co-immunoprecipitation (CoIP)

The cardiomyocytes were collected, and proteins were harvested using a Radioimmunoprecipitation assay (RIPA) lysis buffer (Solarbio). Protein G magnetic beads (Invitrogen) was bound to rabbit polyclonal antibodies against phospho-JNK, and the target antigen (p-JNK) according to manufacturer’s instructions. Subsequently, the co-immunoprecipitation samples were eluted and denatured in 5×SDS-PAGE loading buffer at 95 °C for 5 min, and subsequently analyzed by western blotting with anti-p-JNK anti-P300, anti-PCAF, and anti-H3K9ac antibodies, as described in the section on western blotting. IgG was used as a negative control.

### 2.8. Chromatin immunoprecipitation (ChIP)

After homogenizing the cardiomyocytes, formaldehyde (1%) was added to the samples to cross-link the DNA-protein complexes. ChIP assays were performed using a specific kit (Merck Millipore, Darmstadt, Germany). After crosslinking, the DNA was sheared through ultrasonication, and DNA-protein complex precipitation was performed using monoclonal antibodies (anti-MEF2A, anti-ANP, anti-BNP, anti-β-MHC, anti-P300, and anti-PCAF). The DNA was extracted using a DNA purification kit (Merck Millipore, Darmstadt, Germany). Anti-RNA polymerase II antibody and normal mouse IgG were used as specific antibodies and the negative control, respectively. The ChIP experiments were performed in triplicate. The primer sequences were designed as follows: MEF2A (F) 5’-CAGGTGGTGGCAGTCTTGGA-3’, MEF2A (R) 5’-TGCTTATCCTTTGGGCATTCA-3’; β-MHC (F) 5’-TGAGACGGATGCCATACAGA-3’, β-MHC (R) 5’-GCAGCCTGTGCTTGGTCTT-3’; ANP (F) 5’ -TCCTTGGTGTCTCTCGCTCT-3’, ANP (R) 5’-CGCTGGCTTGCTTGTTGTA-3’; BNP (F) GACAAGAGAGAGCAGGACACCAT, BNP (R) TAAGGAAAAGCAGGAGCAGAATCAT.

### 2.9. Measurements of intracellular Ca^2+^ concentration

Fluo-3/AM (Sigma) was used to probe the intracellular Ca^2+^ concentration. After the cardiomyocytes were treated with indicated concentrations of AA, sp600125 and PE for 48 h, cells were loaded with 10 μmol/L Fluo-3/ AM for 1 h at 37 °C in the dark, followed by analyzed with confocal microscope.

### 2.10. Statistical analysis

All data are shown as the mean ± SD. An LSD-*t* test and one-way analysis of variance (1-way ANOVA) were used to determine statistical significance. SPSS statistical software package version 18.0 (SPSS Inc., Chicago, IL, USA) was used for statistical analysis. A *P*-value < 0.05 was considered statistically significant.

## 3. Results

### 3.1. The activity of HATs in hypertrophic cardiomyocytes induced by PE

To establish a model of PE-induced cardiomyocyte hypertrophy, we first ascertained the optimal PE dose, which was found to be 100 μmol/L, based on the activity of myocardial cells, as well as on the level of *ANP* mRNA expression in cultured cardiomyocytes (Fig. 1A-B). To evaluate PE-induced cardiomyocyte hypertrophy, treated mouse primary myocardial cells were examined by immunofluorescence. PE-treated myocardial cells appeared obviously enlarged compared to control cells (Fig. 1C) and exhibited a substantial increase in cell surface area (Fig. 1D). Some evidence suggests that imbalanced histone acetylation by HATs is involved in PE-induced cardiomyocyte hypertrophy [8]. Hence, we tested the activity of HATs in hypertrophic cardiomyocytes. Colorimetric assays revealed significantly increased HAT activity in hypertrophic cardiomyocytes induced by PE compared to that in normal cells (*P* < 0.05; Fig. 1E). To further confirm the link between histone acetylation and cardiac hypertrophy, the HAT inhibitor AA was administrated to PE-induced hypertrophic myocardial cells. First, the structural formula of AA from Chinese herb extracts was determined (Fig. 1F). Next, the optimal dose of AA was ascertained. To this end, hypertrophic cardiomyocytes were administrated various concentrations of AA (0, 30, 40, 50, 60 μmol/L), based on previous reports [9]. The concentration of 50 μmol/L was selected in accordance with the level of H3K9ac in myocardial cells (Fig. 1G).

**Fig. 1.**
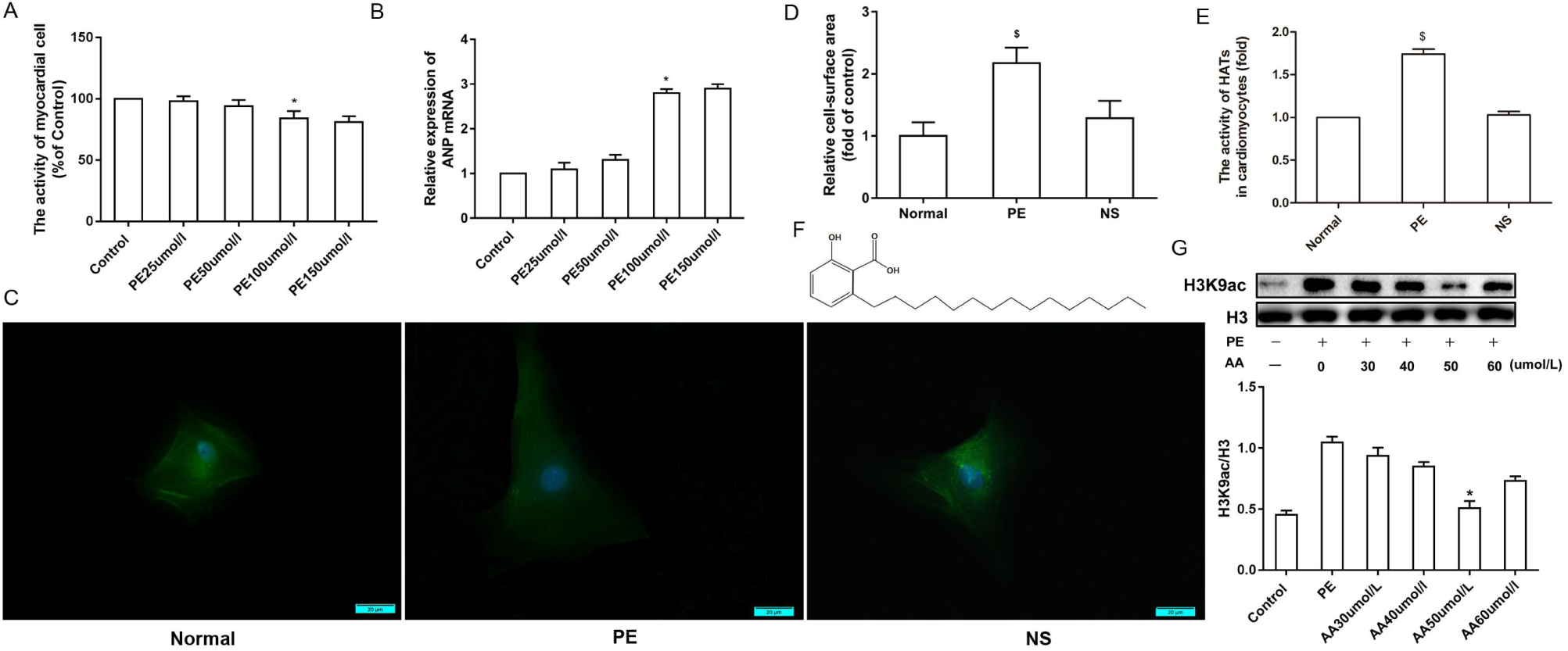
Activity of HATs in hypertrophic cardiomyocytes induced by PE. Different concentrations of PE (25, 50, 100, and 150 μmol/L) were used to determine the optimal PE dose for the induction of cardiomyocyte hypertrophy in neonatal mice. (**A)** The effect of PE on cell viability was accessed using the Cell Counting Kit-8 assay in neonatal mouse cardiomyocytes. (**B)** Real-time PCR analysis of *ANP* mRNA expression after treatment with different concentrations of PE for 48 h. (**C)** The cardiomyocyte cytoplasm was stained with antibodies against α-actin, while the nuclei were stained with DAPI. The scale bars represent 20 μm. (**D)** Quantification of cell surface areas in each treatment group. (**E)** PE (100 μmol/L) increased HAT activity. (**F)** The chemical structure of the HAT inhibitor anacardic acid (AA). (**G)** Different concentrations of AA (30, 40, 50, and 60 μmol/L) were used to identify the optimal dose and 50 μmol/L AA was selected based on the level of H3K9ac. \**P* < 0.05 *vs*. the control group, $*P* < 0.05 *vs*. the normal group (n = 6)

### 3.2. AA attenuates histone H3K9 hyperacetylation via the p-JNK pathway in mouse hypertrophic cardiomyocytes

We next verified whether AA could attenuate the effects of p-JNK signaling on histone H3K9 hyperacetylation in PE-induced hypertrophic cardiomyocytes. First, the optimal concentration of the JNK inhibitor SP600125 (20 μmol/L) was defined by taking into account myocardial cell viability, as assessed by CCK-8 assays (Fig. 2A). The activity of the JNK signaling pathway and the level of H3K9ac were tested by western blotting and/or immunofluorescence. Western blotting showed that in PE-treated cells, the level of p-JNK was significantly increased compared to that in control cells and that both SP600125 and AA attenuated the PE-induced effects, whereas T-JNK was not changed under the same conditions (Fig. 2B). In addition, both immunofluorescence and western blotting showed the occurrence of H3K9 hyperacetylation in PE-treated myocardial cells, whereas the JNK inhibitor SP600125, as well as the HAT inhibitor AA, attenuated PE-induced hyperacetylation (Fig. 2C-E). These data suggested that AA downregulates PE-induced histone H3K9 hyperacetylation via the p-JNK signaling pathway in mouse myocardial cells.

**Fig. 2.**
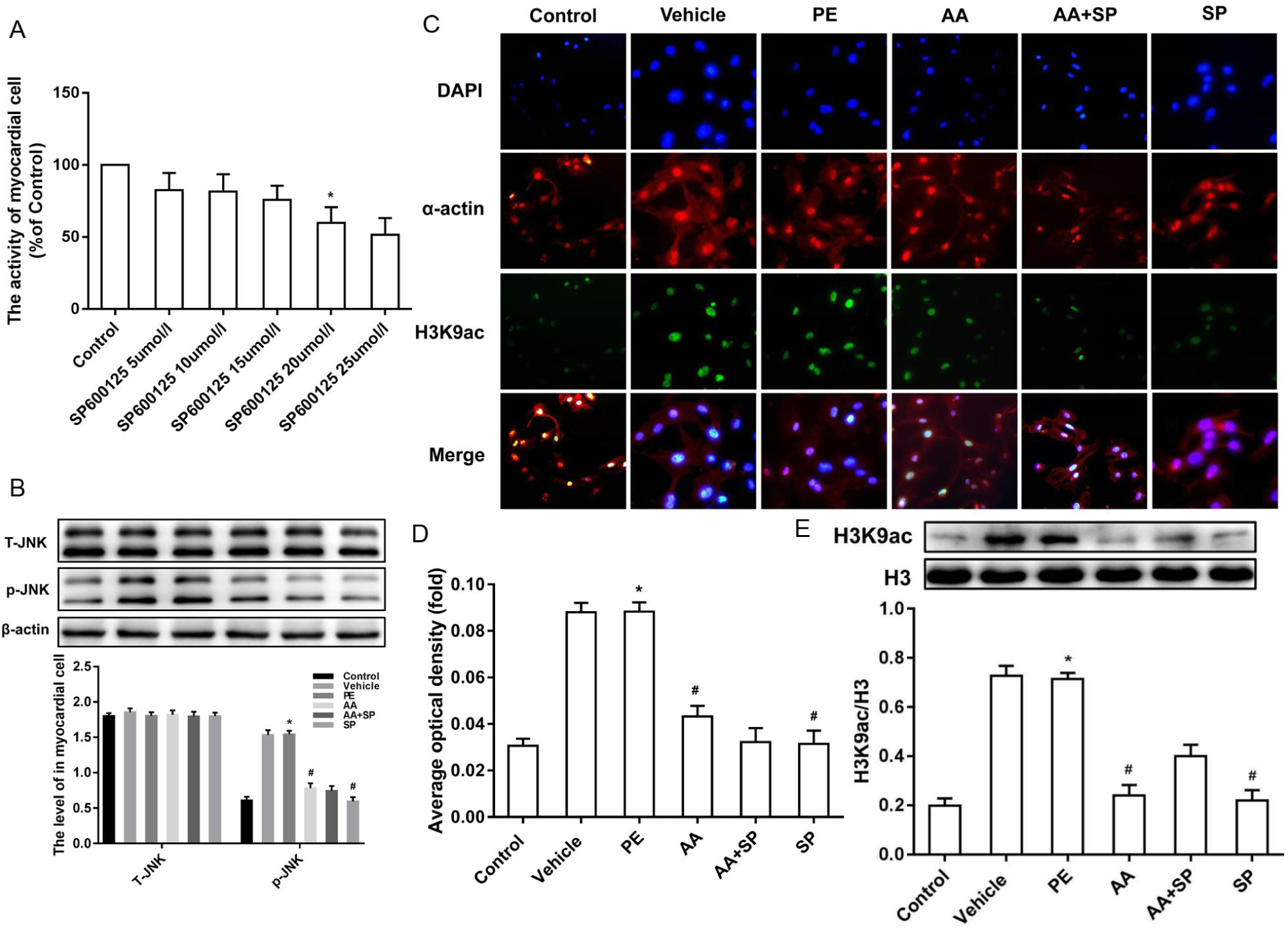
Effects of JNK/MAPK signaling and H3K9ac on mouse hypertrophic cardiomyocytes. **(A)** Effect of different concentrations of the JNK inhibitor SP600125 (5, 10, 15, 20, and 25 μmol/L) on cell viability of neonatal mouse cardiomyocytes. (**B)** Western blotting showing that p-JNK was significantly increased in myocardial cells exposed to phenylephrine (PE); the JNK inhibitor SP600125 decreased p-JNK overexpression in PE-induced hypertrophic cardiomyocytes but had not effect on T-JNK. (**C)** H3K9ac (green fluorescence) and α-actin (red fluorescence) combined with DAPI (blue fluorescence) staining in myocardial cells exposed to six different conditions. Scale bars, 20 μm. (**D)** Average optical density of H3K9ac immunofluorescence in the six groups. All results are representative of at least three independent experiments. (**E)** The level of H3K9ac was significantly increased in PE-induced hypertrophic cardiomyocytes; the HAT inhibitor anacardic acid (AA) attenuated histone H3K9 hyperacetylation in PE-induced hypertrophic cardiomyocytes. \**P* < 0.05 *vs*. the control group, #*P* < 0.05 *vs*. the PE group (n = 3)

### 3.3. p-JNK directly interacts with HATs and alters H3K9 acetylation in primary mouse cardiomyocytes

Evidence supporting an important role for the p-JNK signaling pathway in pathological myocardial cell hypertrophy was previously reported [4]. Our previous study found that alterations in HAT-mediated H3K9ac modifications are involved in pathological myocardial cell hypertrophy [7]. Therefore, we reasoned that the p-JNK signaling pathway could functionally interact with HATs to regulate histone H3K9 acetylation, thus contributing to pathological cardiac hypertrophy. To verify this hypothesis, CoIP experiments were conducted to evaluate the formation of a complex comprising p-JNK and HATs (P300 and PCAF), involved in H3K9 acetylation, and demonstrated the occurrence of such interactions in primary cultured myocardial cells. These data indicated that the p-JNK signaling pathway might have a direct impact on HAT-mediated H3K9 acetylation (Fig. 3A-B).

**Fig. 3.**
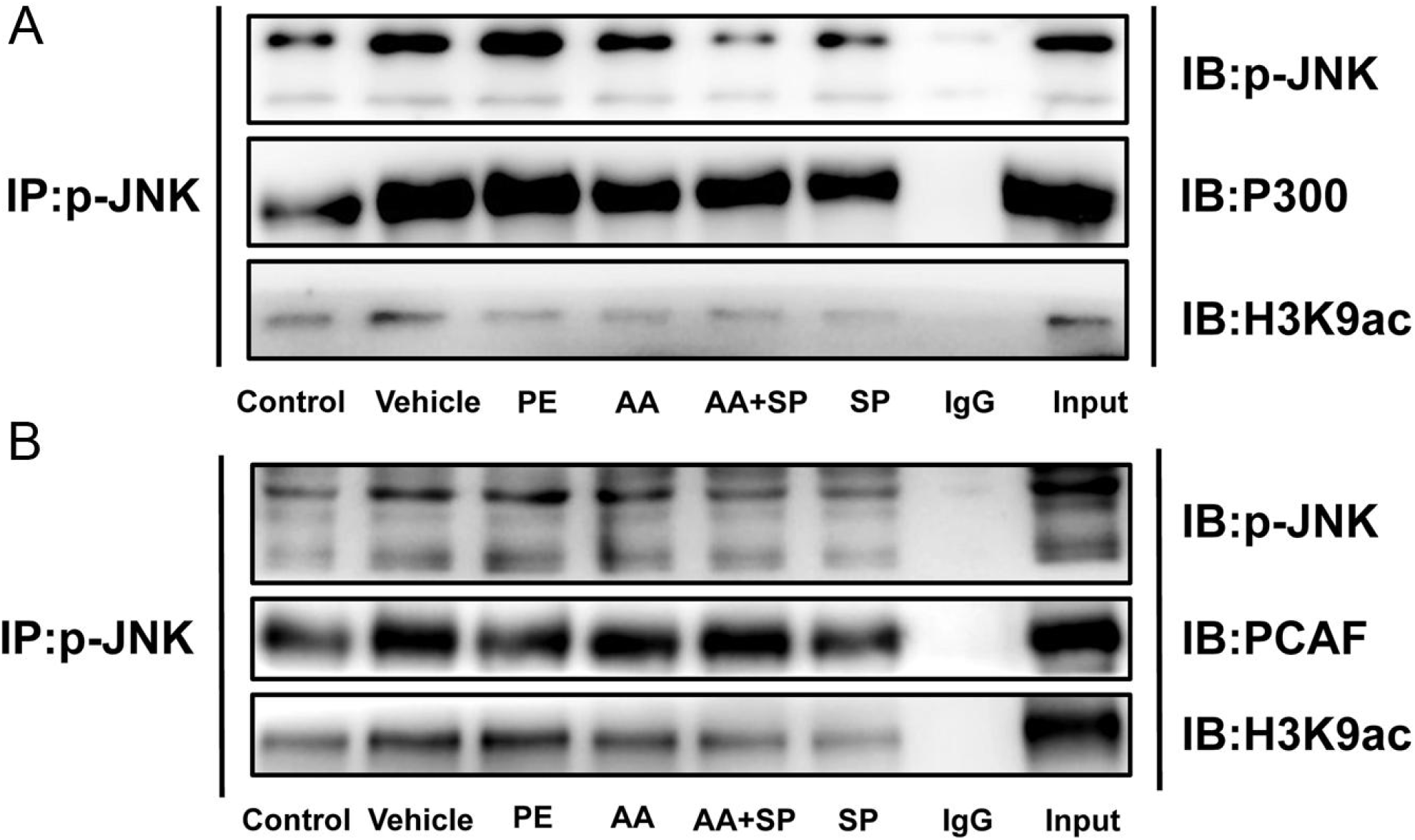
p-JNK directly interacted with HATs and altered H3K9 acetylation in primary mouse cardiomyocytes. **(A and B)** Co-immunoprecipitation (CoIP) of cell lysates from mouse myocardial cells exposed to six different experimental conditions with anti-p-JNK–protein G magnetic beads and immunoblot (IB) with an anti-P300-HAT, anti-PCAF-HAT, anti-H3K9ac-HAT, or anti-p-JNK antibody for the evaluation of protein expression. Input: positive control; IgG: negative control

### 3.4. AA attenuates the overexpression of p300-HAT and PCAF-HAT induced by PE via the p-JNK pathway

HATs and histone deacetylases (HDACs) are involved in the regulation of histone acetylation. Our previous study suggested that P300-HAT and PCAF-HAT play a critical role in pathological cardiac hypertrophy by affecting histone acetylation. Thus, we performed immunofluorescence and western blotting to analyze the expression of P300-HAT. Treatment with PE induced an obvious increase in P300-HAT expression, whereas exposure to both AA and the JNK inhibitor SP600125 attenuated P300-HAT overexpression in PE-treated mouse primary myocardial cells (Fig. 4A-C). The expression of PCAF-HAT was also assayed by immunofluorescence and western blotting in the same samples. The data showed that PCAF-HAT expression was significantly increased in PE-treated cardiomyocytes compared to that in control cells. However, the HAT inhibitor AA or the JNK inhibitor SP600125 partially prevented PE-the induction of PCAF-HAT overexpression (Fig. 4D-F).

**Fig. 4.**
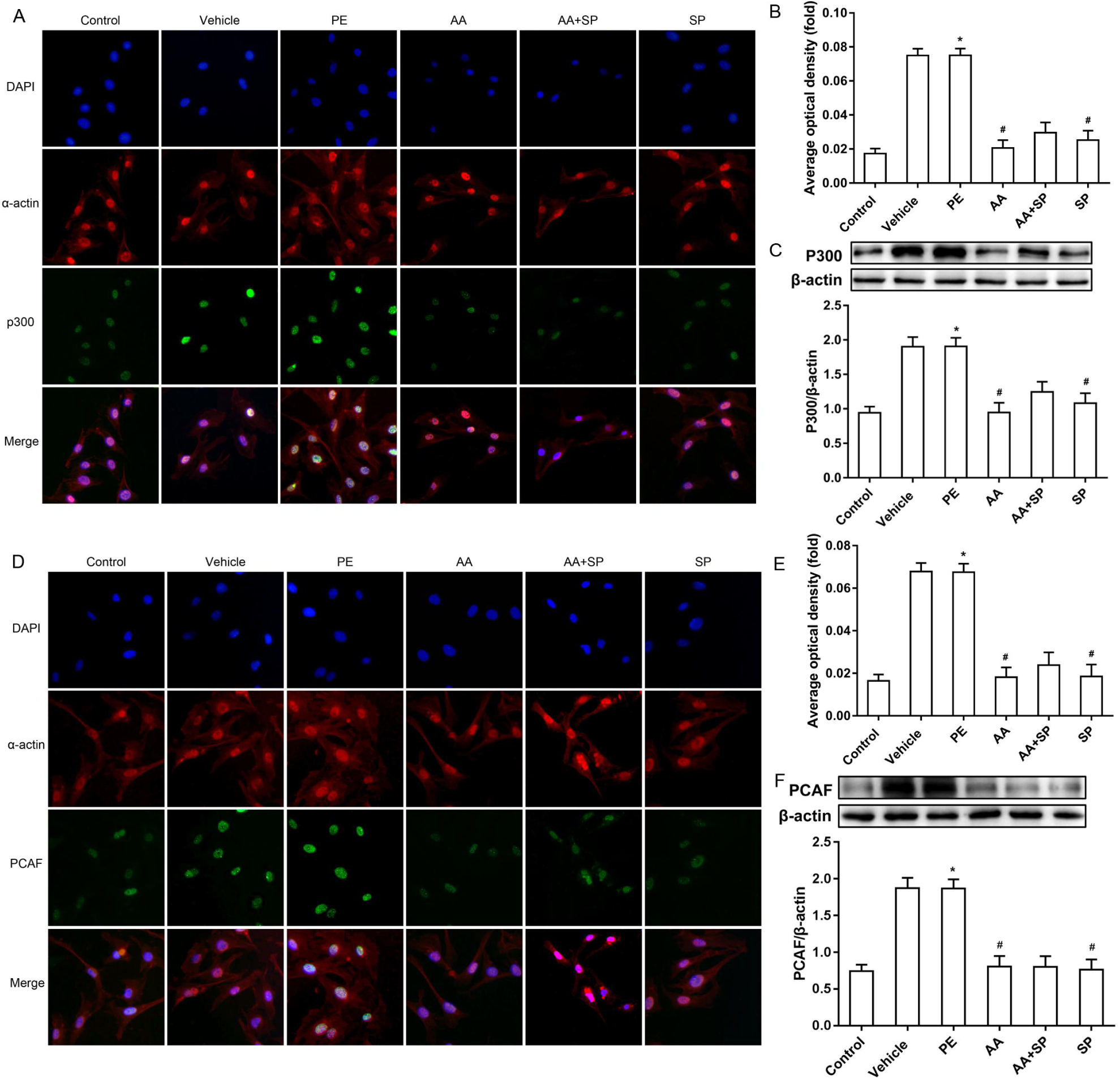
Effects of AA and the JNK inhibitor SP600125 on the expression of HATs in PE-induced hypertrophic cardiomyocytes. **(A)** P300 (green fluorescence) and α-actin (red fluorescence) combined with DAPI (blue fluorescence) staining of myocardial cells exposed to six different conditions. Scale bars, 20 μm. (**B)** Average optical density of P300 immunofluorescence in the six groups. (**C)** The levels of P300-HAT (histone acetyltransferase) significantly increased in the hypertrophic cardiomyocytes induced by PE; the HAT inhibitor AA and the JNK inhibitor SP600125 prevented this effect. (**D)** PCAF (green fluorescence) and α-actin (red fluorescence) combined with DAPI (blue fluorescence) staining of myocardial cells exposed to six different conditions. Scale bars, 20 μm. (**E)** Average optical density of PCAF immunofluorescence in the six groups. (**F)** The level of PCAF-HAT was increased significantly in hypertrophic cardiomyocytes induced by PE compared to that in control cells. The HAT inhibitor AA or the JNK inhibitor SP600125 normalized the expression of PCAF-HAT in PE-induced hypertrophic cardiomyocytes. \**P* < 0.05 *vs*. the control group, #*P* < 0.05 *vs*. the PE group (n = 6)

### 3.5. AA attenuates the PE-induced and p-JNK signaling-dependent overexpression of cardiac hypertrophy-related genes

The effect of HAT-mediated regulation of the cardiac nuclear transcription factor MEF2A on the remodeling of pathological cardiomyocytes was evaluated by ChIP-PCR. These results showed that P300 and PCAF could bind the *MEF2A* promoter, suggesting their involvement in the regulation of this transcription factor (Fig. 5A). Some evidence suggests that *MEF2A* is a critical factor in pathological cardiac hypertrophy [10,11]. Hence, the transcriptional level of *MEF2A* was examined using qRT-PCR, showing that it was significantly upregulated in PE-treated cells compared to levels in control cells. Meanwhile, the HAT inhibitor AA or the JNK inhibitor SP600125 suppressed PE-induced *MEF2A* overexpression in primary cultured myocardial cells (Fig. 5B). In addition, ChIP-PCR showed that MEF2A could bind the promoters of cardiac hypertrophy biomarker genes such as *ANP, BNP*, and *β-MHC*, suggesting its role in the transcriptional regulation of these genes (Fig. 5C). In addition, the protein expression of ANP, BNP, and β-MHC was assayed by western blotting. The levels of these proteins were obviously increased in PE-treated cells compared to those in control cells, whereas this effect was attenuated by both AA and SP600125 (Fig. 5D-F).

**Fig. 5.**
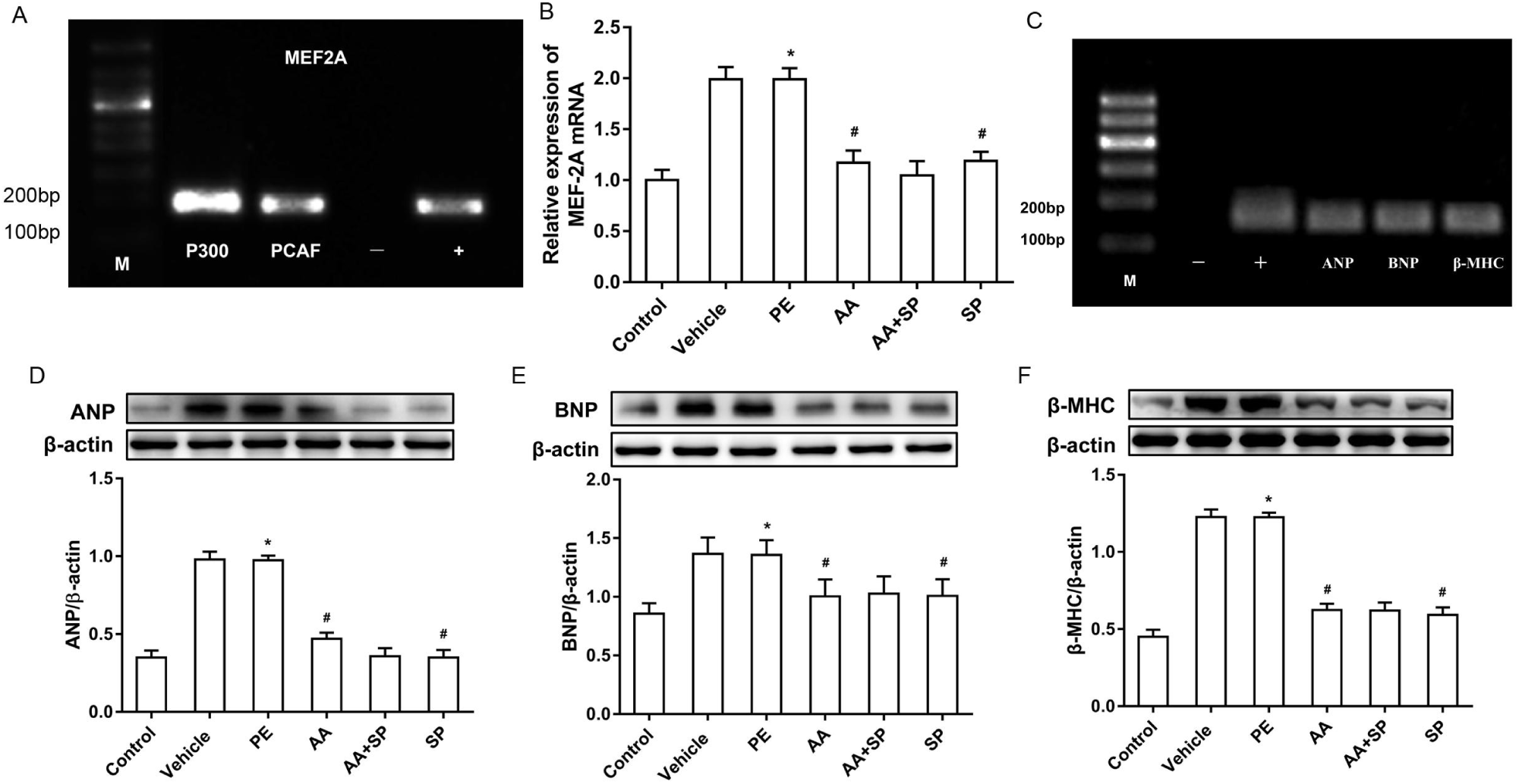
Effects of AA and the JNK inhibitor SP600125 on the expression of cardiac hypertrophy-related genes in PE-induced hypertrophic cardiomyocytes. **(A)** Chromatin immunoprecipitation (ChIP)-PCR results demonstrating the binding of P300 and PCAF to the *MEF2A* promoter. (-) negative control, amplification of DNA fragments after precipitation with normal mouse IgG; (+) positive control, amplification of DNA fragments after precipitation with anti-RNA polymerase II antibody. (**B)** qRT-PCR results showing that the mRNA expression of *MEF2A* was higher in PE-induced hypertrophic cardiomyocytes than in control cardiomyocytes, whereas histone acetyltransferase (HAT) and JNK inhibition with AA and SP600125, respectively, prevented *MEF2A* overexpression in these cells. (**C)** ChIP-PCR demonstrated that *MEF2A* could bind the promoters of atrial natriuretic peptide (*ANP*), brain natriuretic peptide (*BNP*), and beta-myosin heavy chain (*β-MHC*). Input: positive control, IgG: negative control. (**D, E, and F)** Western blotting showing that expression of the cardiac hypertrophy-related proteins ANP, BNP, and β-MHC was higher in the PE group than in the control group, whereas the HAT inhibitor AA or the JNK inhibitor SP600125 counteracted this effect. \**P* < 0.05 *vs*. the control group, #*P* < 0.05 *vs*. the PE group (n = 6)

### 3.6. AA attenuates cardiomyocyte hypertrophy and improves cardiac function via the p-JNK signaling pathway

We then investigated the impact of p-JNK-mediated alterations in histone acetylation on PE-induced cardiac hypertrophy. In particular, the surface area of cardiomyocytes was evaluated by immunofluorescence. In PE-treated cardiomyocytes, the surface area was significantly increased compared to that in control cells, whereas both HAT and JNK inhibition reduced this increase (Fig. 6A-B). It is known that the intracellular Ca^2+^ concentration is an important indicator of myocardial systolic function [12]. Hence, this parameter was assayed by confocal laser scanning microscopy. In PE-treated cardiomyocytes, intracellular Ca^2+^ was clearly increased compared to that in control cells, whereas both HAT inhibition by AA and JNK inhibition by SP600125 reduced this effect (Fig. 6C-E).

**Fig. 6.**
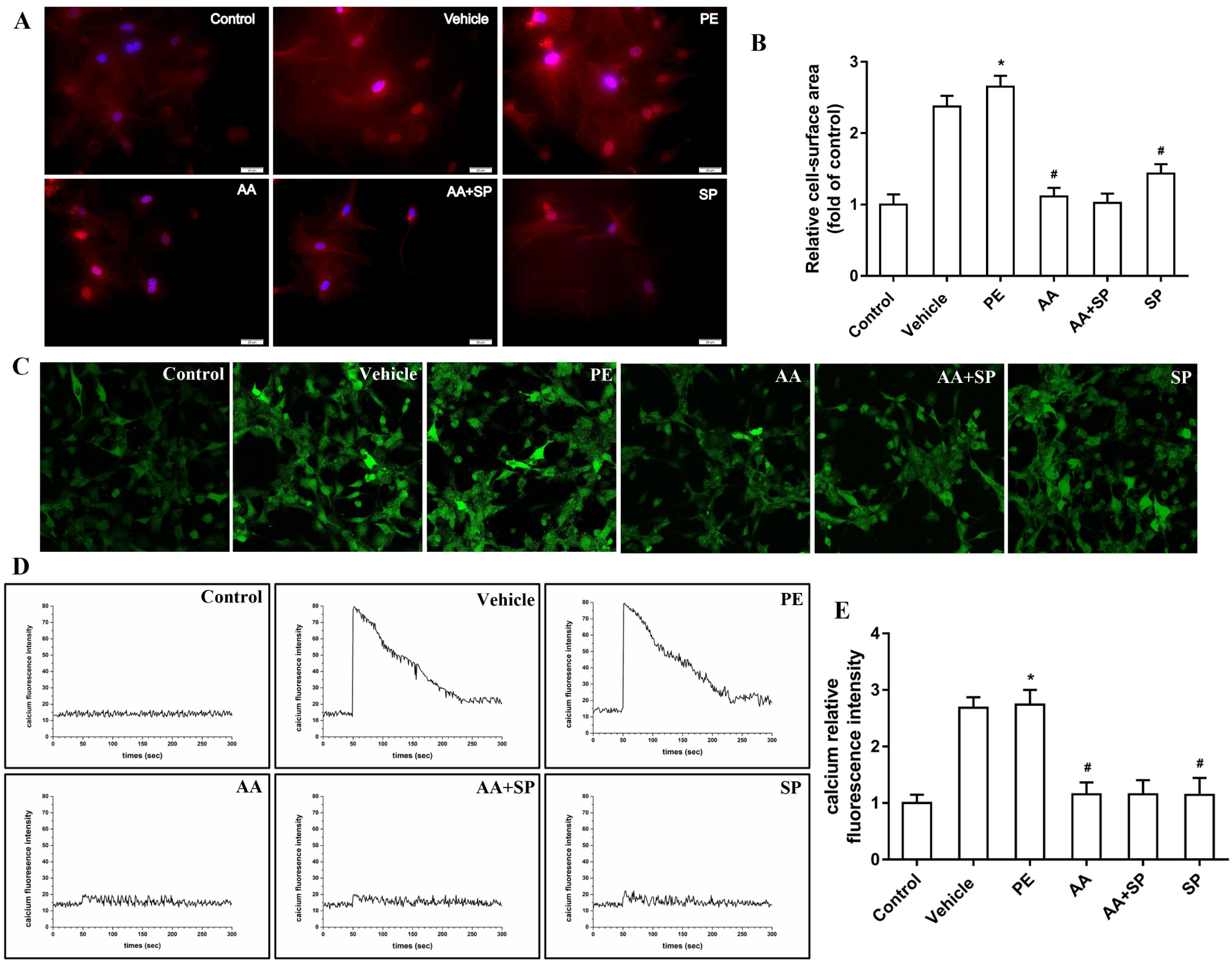
The HAT inhibitor AA and the JNK inhibitor SP600125 attenuate cardiomyocyte hypertrophy induced by PE. **(A)** The cell surface area was measured by immunofluorescence staining to demonstrate hypertrophic responses in cardiomyocytes. (**B)** Measurement of the cell surface area of myocardial cells for the quantification of cell size. (**C)** Determination of intracellular Ca^2+^ concentrations using the calcium fluorescent probe Flou-3/AM and laser scanning confocal microscopy. (**D)** Dynamic fluorescence curve of intracellular Ca^2+^ in myocardial cells. (**E)** Comparison of intracellular Ca^2+^ fluorescence intensity in myocardial cells among the six groups of PE-induced hypertrophic cardiomyocytes. \**P* < 0.05 *vs*. the control group, #*P* < 0.05 *vs*. the PE group (n = 6). Scale bars, 20 μm

## 4. Discussion

Epigenetic regulation of gene expression contributes to the pathogenesis of various human diseases [13]. Evidence is mounting that alterations to multiple signaling pathways might result in epigenetic changes that, in turn, regulate the transcription of biomarker genes of cardiac hypertrophy [14]. In this study, we demonstrated that AA-mediated protection from cardiac hypertrophy occurs via the modification of histone acetylation, which in turn was found to be regulated by the JNK/MAPK signaling pathway, as illustrated in (Fig. 7). These findings shed new light on the therapeutic effects of AA on pathological cardiac hypertrophy.

**Fig. 7.**
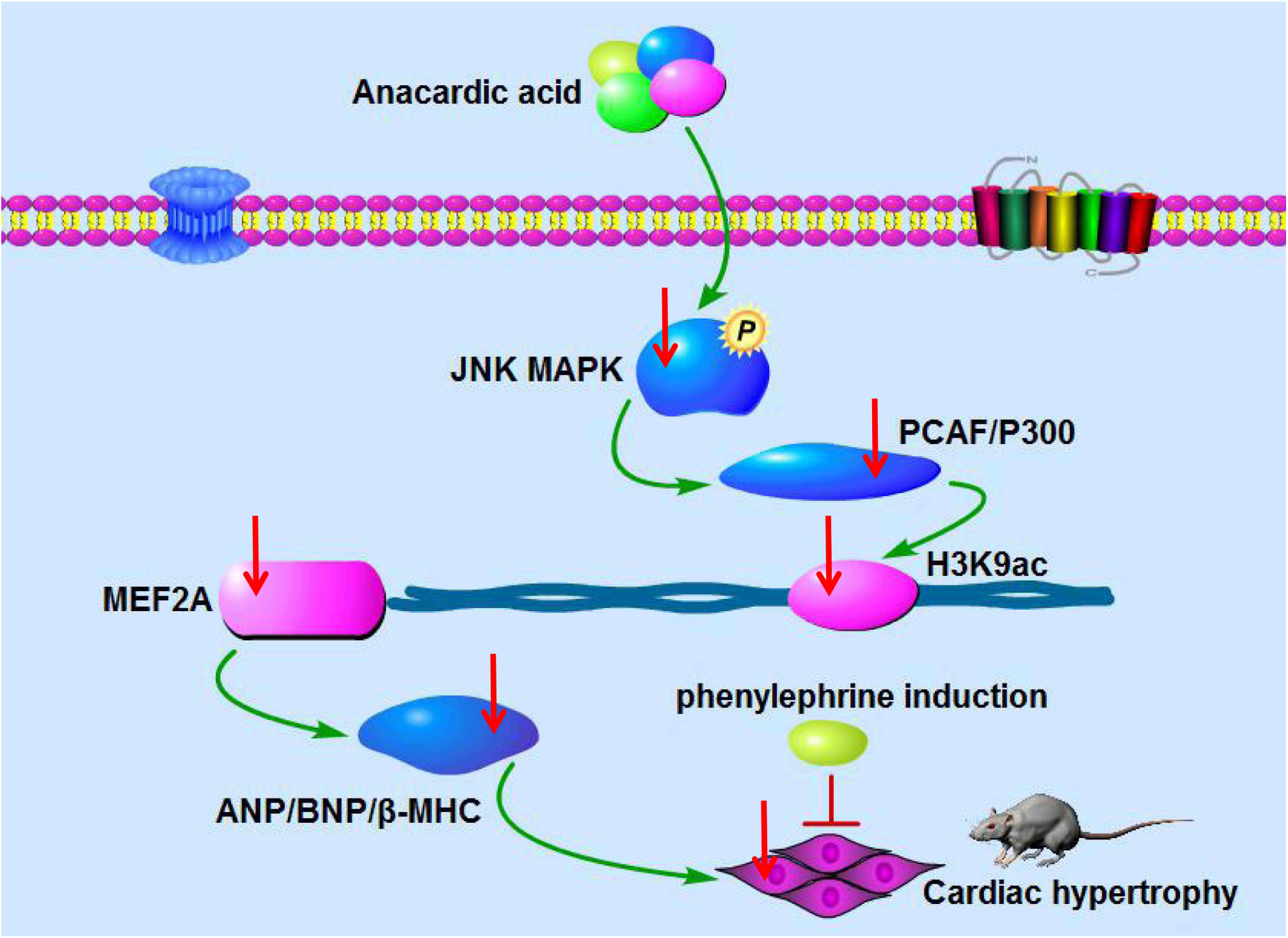
Schematic representation of the possible mechanism through which AA attenuates PE-induced cardiomyocyte hypertrophy. AA could ameliorate cardiac hypertrophy by regulating JNK signaling-induced histone acetylation

Cardiac hypertrophy is a pathological reaction induced by a variety of factors and is characterized by an increase in cardiomyocyte volume and protein synthesis [15,16]. This condition initially develops as a compensatory ventricular hypertrophy. If not treated properly, this adaptive process might gradually develop into arrhythmia, heart failure, and even sudden cardiac death due to decompensation [17]. However, the events underlying the conversion from adaptive to maladaptive hypertrophy are yet to be defined. Therefore, extensive efforts are being made to develop more effective treatment measures. Recently, numerous studies have shown that several intracellular signaling pathways might affect the progression of cardiac hypertrophy [18,19]. Notably, it has recently been shown that the hyperactivation of MAPK signaling plays a crucial role in the development of cardiac hypertrophy [20]. JNK is a member of the MAPK pathway and its altered expression is associated with cardiac hypertrophy [21]. A previous study indicated that the JNK/MAPK signaling pathway is activated during cardiomyocyte hypertrophy and that its inhibition alleviates this condition [22]. Moreover, it was reported that JNK-interacting protein 3 (JIP3) knockdown suppresses JNK signaling and attenuates cardiac hypertrophy [23]. In addition, Harsha Raj et al. found that AA induces apoptosis in cancer cells by regulating MAPK signaling [24]. In recent years, increasing lines of evidence have suggested that PE is an important neurohumoral factor closely involved in the pathogenesis of cardiac hypertrophy [25]. Our previous studies demonstrated that AA attenuates PE-induced cardiac hypertrophy [7]. However, whether JNK/MAPK signaling plays a role in this effect was not clear.

One study reported that JNK inhibition via SP600125 attenuates cardiac hypertrophy induced by pressure overload [26]. Our experiments showed that the level of p-JNK protein was significantly increased in PE-treated cardiomyocytes compared to that in control cells, whereas the JNK inhibitor SP600125 prevented this overexpression. Moreover, our data indicated the occurrence of H3K9 hyperacetylation in PE-treated cardiomyocytes and that the HAT inhibitor AA attenuates this effect. A previous study from our laboratory showed that imbalanced HAT-mediated H3K9 acetylation is involved in pathological cardiac hypertrophy [7]. Hence, we reasoned that p-JNK could interact with HATs, thereby affecting H3K9ac and contributing to cardiac hypertrophy. CoIP experiments were performed to verify this hypothesis, indicating that p-JNK directly interacts with HATs (P300-HAT and PCAF-HAT). Previous evidence suggested that altered histone acetylation by HATs is implicated in PE-induced cardiomyocyte hypertrophy [27]. Our experiments showed a significant increase in HAT activity in hypertrophic cardiomyocytes induced by PE. Recently, the role of HATs and HDACs in the regulation of histone acetylation has been confirmed [28]. We previously showed that P300-HAT and PCAF-HAT play a critical role in pathological cardiac hypertrophy and that this effect depends on their ability to modify histone acetylation [7]. Our data indicated that PE significantly increases P300-HAT and PCAF-HAT expression in hypertrophic cardiomyocytes, whereas both HAT and JNK blockade were found to attenuate these effects. Many studies have shown that the cardiac nuclear transcription factor MEF2A is a critical regulator of pathological cardiac hypertrophy [29,30]. Interestingly, substantial *MEF2A* overexpression was observed by qRT-PCR in PE-treated cardiomyocytes. Notably, the HAT inhibitor AA or the JNK inhibitor SP600125 partially prevented this PE-induced *MEF2A* upregulation. Furthermore, we directly addressed the relationship between *MEF2A* and HATs. In particular, ChIP-PCR indicated that P300-HAT and PCAF-HAT were involved in the regulation of *MEF2A* expression. In addition, western blotting analysis showed that protein levels of ANP, BNP, and β-MHC were increased in PE-treated cardiomyocytes compared to those in control cells and that HAT or JNK inhibition could attenuate these effects. Furthermore, ChIP-PCR suggested that MEF2A regulates the expression of *ANP, BNP*, and *β-MHC*. Moreover, immunofluorescence experiments showed that the surface area of PE-exposed cardiomyocytes was significantly increased compared to that in controls and that HAT or JNK inhibition could partially prevent this. Ca^2+^ plays a key regulatory role in the metabolic activity and systolic function of cardiomyocytes [12]. Therefore, we also measured the intracellular Ca^2+^ concentration with Fluo-3/AM. In this regard, laser confocal microscopy showed that in PE-treated cardiomyocytes, the intracellular Ca^2+^ concentration was increased compared to that in controls and that HAT or JNK inhibition could reduce the PE-induced increase in intracellular Ca^2+^. Therefore, AA alleviates PE-induced cardiomyocyte hypertrophy and improves cardiomyocyte function by modifying histone acetylation via the JNK/MAPK signaling pathway.

Our findings provide new insight into the treatment of cardiac hypertrophy. Nonetheless, the participation of other members of the MAPK protein family in the AA-induced attenuation of cardiac hypertrophy cannot be ruled out. Future studies are urgently needed to explore this issue.

## Abbreviations

AA: anacardic acid
ANP: atrial natriuretic peptide
BNP: brain natriuretic peptide
CoIP: Co-immunoprecipitation
HAT: histone acetyltransferase
HDAC: histone deacetylase
H3K9: histone H3 lysine 9
H3K9ac: acetylated lysine 9 on histone H3
JNK: c-Jun N-terminal Kinase
PCAF: P300/CBP-associated factor
PE: phenylephrine.

## Acknowledgments

Conceived and designed the experiments: BHP and CP. Performed the experiments: BHP, XML, and LXH. Analyzed the data: CP. Provided reagents, materials, and analysis tools: QM, XH, and HTZ. Wrote the original draft: BHP and CP. This study was supported by the National Natural Science Foundation of China (grant number: 81560040), the Program of Science and Technology Department of Guizhou Province of China (grant number: [2016]1177), and the Doctoral Startup Foundation of the Affiliated Hospital of Zunyi Medical University (grant number: 2015-4).

